# An evaluation of the accuracy and speed of metagenome analysis tools

**DOI:** 10.1101/017830

**Authors:** Stinus Lindgreen, Karen L. Adair, Paul P. Gardner

## Abstract

Metagenome studies are becoming increasingly widespread, yielding important insights into microbial communities covering diverse environments from terrestrial and aquatic ecosystems to human skin and gut. With the advent of high-throughput sequencing platforms, the use of large scale shotgun sequencing approaches is now commonplace. However, a thorough independent benchmark comparing state-of-the-art metagenome analysis tools is lacking. Here we present a benchmark where the most widely used tools are tested on complex, realistic data sets. Our results clearly show that the most widely used tools are not necessarily the most accurate, that the most accurate tool is not necessarily the most time consuming, and that there is a high degree of variability between available tools. These findings are important as the conclusions of any metagenomics study are affected by errors in the predicted community composition and functional capacity. Data sets and results are freely available from http://www.ucbioinformatics.org/metabenchmark.html

---

With the advent of second-generation sequencing platforms such as 454^1^ and Illumina^2^, the ease with which any research group can generate gigabases of high-quality sequence data at comparatively low cost has had a major impact on biological research. One field of research that has grown immensely in recent years is metagenomics^3^ - the investigation of full communities to address questions about the composition, diversity and functioning of complex microbial ecosystems. Where the field was initially focused on amplicon sequencing of specific marker genes, today it is commonplace to do high-throughput shotgun sequencing of total DNA in a sample. The result is large, complex data sets that can be used to investigate both taxonomic composition and, potentially, functional capacity of a sample.

Metagenomic studies have shed light on novel aspects of biology. For instance, studies of the human microbiome have shown possible connections between the gut microbiome and diseases as diverse as diabetes^4^, depression^5^ and rheumatoid arthritis^6^. In ecology, metagenomics make it possible to investigate complex communities in e.g. soil^7^, glaciers^8^ and air^9^. Some groups are looking at ancient communities and tracing community changes over time by investigating the metagenomes in coprolites^10^, teeth^11^ and elsewhere^12^. In metatranscriptomics, the expressed RNAs are investigated^13–15^, potentially making it possible to infer transcription patterns directly, but as this field is still in its infancy due to technical challenges we focus on metagenomics in this paper.

The results of any metagenomics study relies on computational tools that can analyze large data sets, and extract useful and correct information about the community under investigation. Fortunately, a number of tools have been developed to investigate the taxonomic composition of metagenomes and, in some cases, also shed light on the functional composition of the community. These tools can broadly be separated in two groups - those using all available sequences for each data set (e.g. CLARK^16^, Genometa ^17^, GOTTCHA^18^, Kraken ^19^, LMAT ^20^, MEGAN ^21,22^, MG-RAST ^23^, the One Codex webserver, taxator-tk ^24^) and those focusing on a set of marker genes (e.g MetaPhlAn ^25^, MetaPhyler ^26^, mOTU ^27^, QIIME ^28^).

A previous benchmark used synthetic communities by pooling genomic DNA in order to primarily investigate different sequencing approaches^29^. However, an independent, in-depth benchmark of analysis tools is lacking. Here we present what is to our knowledge the first unbiased, comprehensive benchmark of metagenome analysis tools in which the authors are not involved in any of the tools tested. We use a set of sequences that were sampled from a known taxonomic distribution, these include genomic sequences from characterised taxa, simulated relatives of varying evolutionary distances using phylogenetic modelling, and permuted sequences that serve as a negative control. These are realistic synthetic metagenomes that capture many of the complexities encountered in real sequencing studiess. We evaluate 14 tools in order to investigate how well they perform both in terms of taxonomy and, where available, function. This comprehensive benchmark sheds light on the performance of widely used tools and can help researchers decide what tools to use in their metagenomics studies, potentially impacting the results and conclusions of any study.

## Results

### Overall performance

The key results are listed in Table 1. The run time varies between tools, ranging from minutes (OneCodex, QIIME) and hours (e.g. mOTU, Kraken) to several days (e.g. MetaPhyler, EBI webserver). However, it should be noted that the input to QIIME is much smaller than the full data sets being analyzed by the other tools as it only contained predicted 16S rRNA sequences. The run time experienced by a user when using a webserver (EBI, MG-RAST, OneCodex) may depend heavily on a number of factors such as current load, software upgrades, and priority of the job.

**Table 1:**
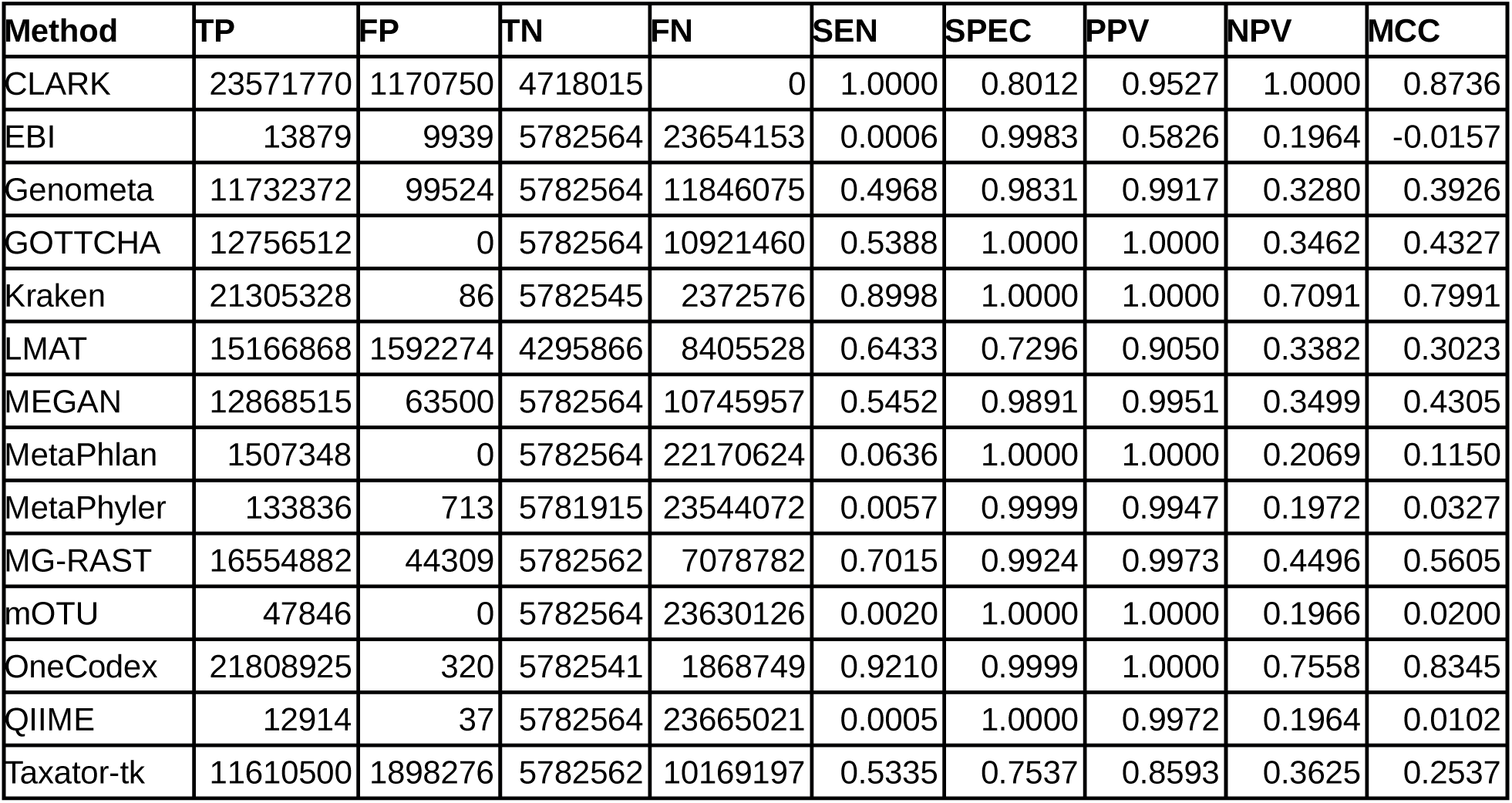
*Performance of the selected tools. Where applicable, the best performance in each category is highlighted in bold. Fraction: average fraction of all reads that the tool mapped. Shuffled: average number of shuffled reads mapped. False positives: fraction of mapped reads assigned to non-existing phyla. Run time: CPU time in minutes per metagenome (where applicable). Correlation: the average Pearson correlation coefficient between predicted and known relative abundances of phyla in the data sets. GOTTCHA failed on setB3 so the average of sets B1 and B2 is used instead.*

The 6 data sets were all designed so that 80% of the reads could ideally be mapped to their corresponding genomes of origin (70% bacteria and archaea, 5% *in silico* evolved bacterial genomes, 5% Eukaryote), and 20% of the reads were sampled from shuffled genomes that should not map. Depending on whether the database used contains Eukaryote genomes or not, each tool should be able to map up to 75%-80% of the reads. All the read counts reported below are averages over the 6 data sets. In total, 29,597,475 read pairs were generated for each data set in this benchmark.

Note, however, that the fraction of reads mapped is not in itself a metric of quality since some tools rely on smaller sets of marker genes. As with the run times, the actual fraction of reads mapped varies greatly between tools. MetaPhlAn, MetaPhyler and mOTU all rely on sets of marker genes, and the EBI webserver uses only predicted rRNA reads, and therefore these four methods map the fewest reads (from 0.08% to appr. 5%). CLARK, Kraken and OneCodex all map >70%, whereas Genometa, MEGAN and Taxator-tk all map around 40%, and LMAT, MGRAST and QIIME map up to 60% of reads. Interestingly, the number of reads analyzed is not reflected in the run time (e.g. Kraken analyzes many more reads than MetaPhyler but runs much faster).

The 5,919,504 (20%) read pairs from shuffled bacterial genomes should not be assigned to any taxa, and indeed the majority of tools assign very few shuffled reads to phyla. Of the 14 methods tested, four map no shuffled reads (EBI, Genometa, MetaPhlAn, QIIME), and three tools map fewer than 30 shuffled reads (Kraken, MG-RAST, Taxator-tk, OneCodex). MetaPhyler maps more than 600 reads, CLARK maps more than 340,000 reads, whereas LMAT maps a large number of shuffled reads to the database (1,486,699 reads). For GOTTCHA, MEGAN and mOTU this information was not readily available.

Another important measure is the occurrence of false positives, i.e. how often a tool predicts the presence of phyla that were not included in the data sets. In general, the tools are consistently specific in that they do not mistakenly predict phyla that aren’t there. Of the reads being mapped, most tools assign less than 1% to phyla other than those included in the simulation. However, there are two outliers with Taxator-tk assigning around 14% of mapped reads to phyla not represented in the simulated data sets, and the EBI server assigning almost 42% of the mapped reads to “Other” bacteria and archaea although not to any specific phyla.

### Taxonomic analysis

We evaluate the performance of each tool by looking at the assignments at both the level of phylum and genus. The data sets contain sequences from 17 phyla (including a single combined “Eukaryote phylum”) and 417 different genera (including 7 Eukaryote genera), see Supplementary Tables 1 and 4 for details. Performance was analyzed both with and without Eukaryotes, but in the following only the results excluding Eukaryotes are described as the relative performance was not significantly affected by this. See Supplementary Figures 3 and 4 (phyla) and 5 and 6 (genera) for performance including all groups. We expect the methods to perform best at identifying phyla, but it is of interest to evaluate the performance at better resolution (genus level) as well as investigate how well the performance correlates between these two levels.

For each tool, the predicted relative abundance of each phylum and genus was compared to the known abundance (see Methods for details on the composition of data sets). The predicted abundance differed significantly from the known abundance (Student’s t-test on every single genus or phylum; all P<0.05) in almost every case (see Supplementary Figure 1 and Supplementary Table 1 for details). Only four tools include Eukaryotes (GOTTCHA, MG-RAST, MetaPhlAn, OneCodex) and all significantly underrepresent the Eukaryote component. However, in many cases the Eukaryote databases only include a small number of organisms, such as selected fungi. Overall, there was a large variation in the predicted relative abundance of phyla. Interestingly, it is not only the rare phyla that show large variation in predicted abundance, also two of the most abundant phyla - Acidobacteria and Proteobacteria - showed large variation between tools.

**Figure 1:**
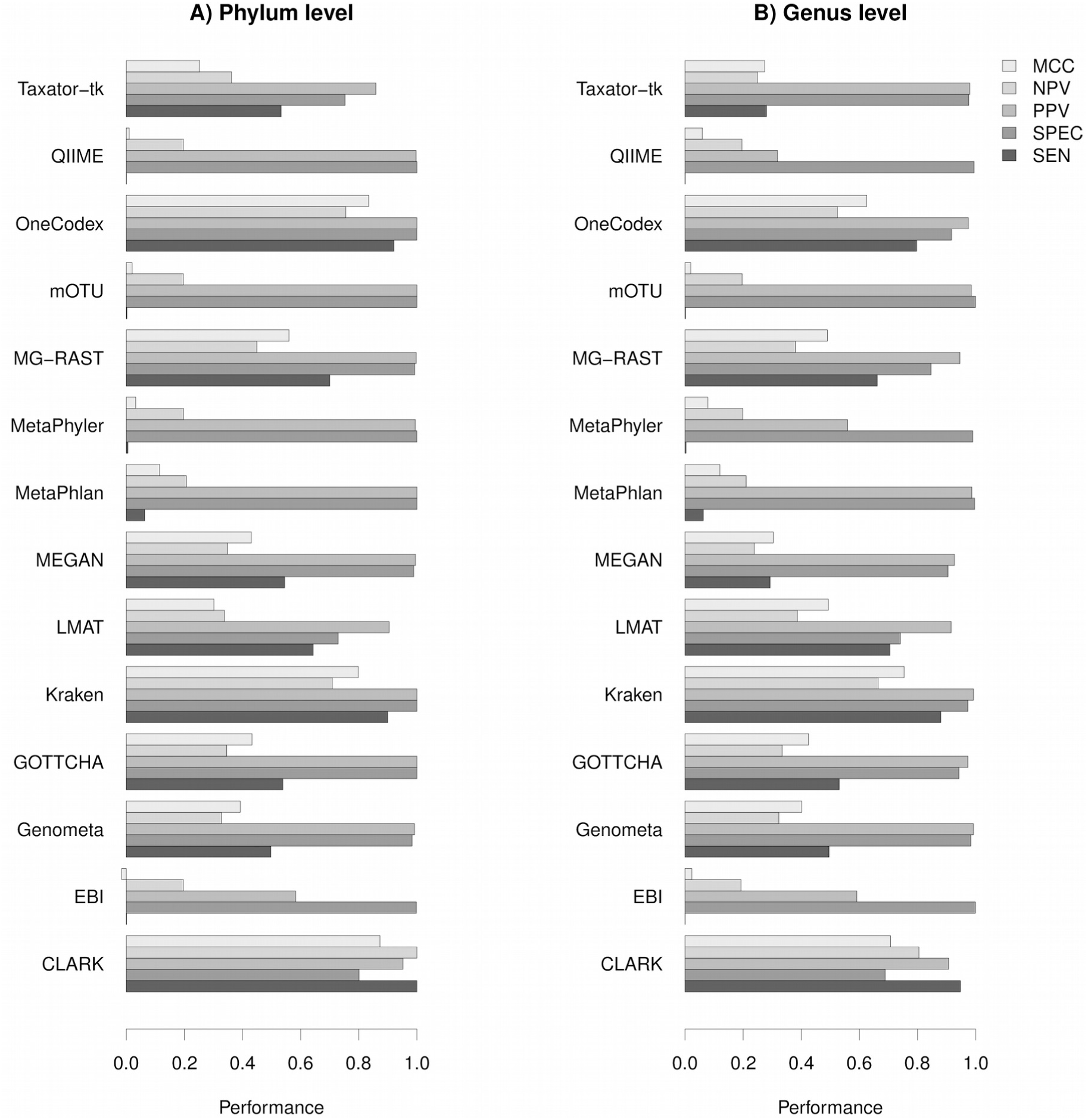
*Plot of performance metrics for all methods included in this benchmark. The closer each metric is to 1, the more accurate the method. The values are based on classifying reads mapped at the level of phylum (A) or genus (B).*

In Table 2, we show the number of true positives (TP), false positives (FP), true negatives (TN) and false negatives (FN) as well as sensitivity (SEN), specificity (SPEC), positive predictive value (PPV), negative predictive value (NPV) and Matthew’s Correlation Coefficient (MCC), see Supplementary Table 1 for details. The performance for each method given these metrics is illustrated in Figure 1, where values close to 1 are best. Overall, the methods perform well in terms of specificity at both the level of phylum (Figure 1A) and genus (Figure 1B). It should be noted that the sensitivity scores for QIIME, EBI, mOTU, MetaPhyler and MetaPhlan are artificially low as all these methods use custom databases that only contain specific marker sequences. Thus, we would not expect to map all reads using these tools, and it is important to also consider the other quality metrics presented in this paper. MCC measures the balance between sensitivity and specificity and can be used as a combined performance metric. Using this metric, we see a strong positive *(r=0.96)* and highly significant (P<10^−7^) Pearson correlation between the phylum and genus level predictions (see Table 3 and 4 for this and other correlations).

**Table 2:**
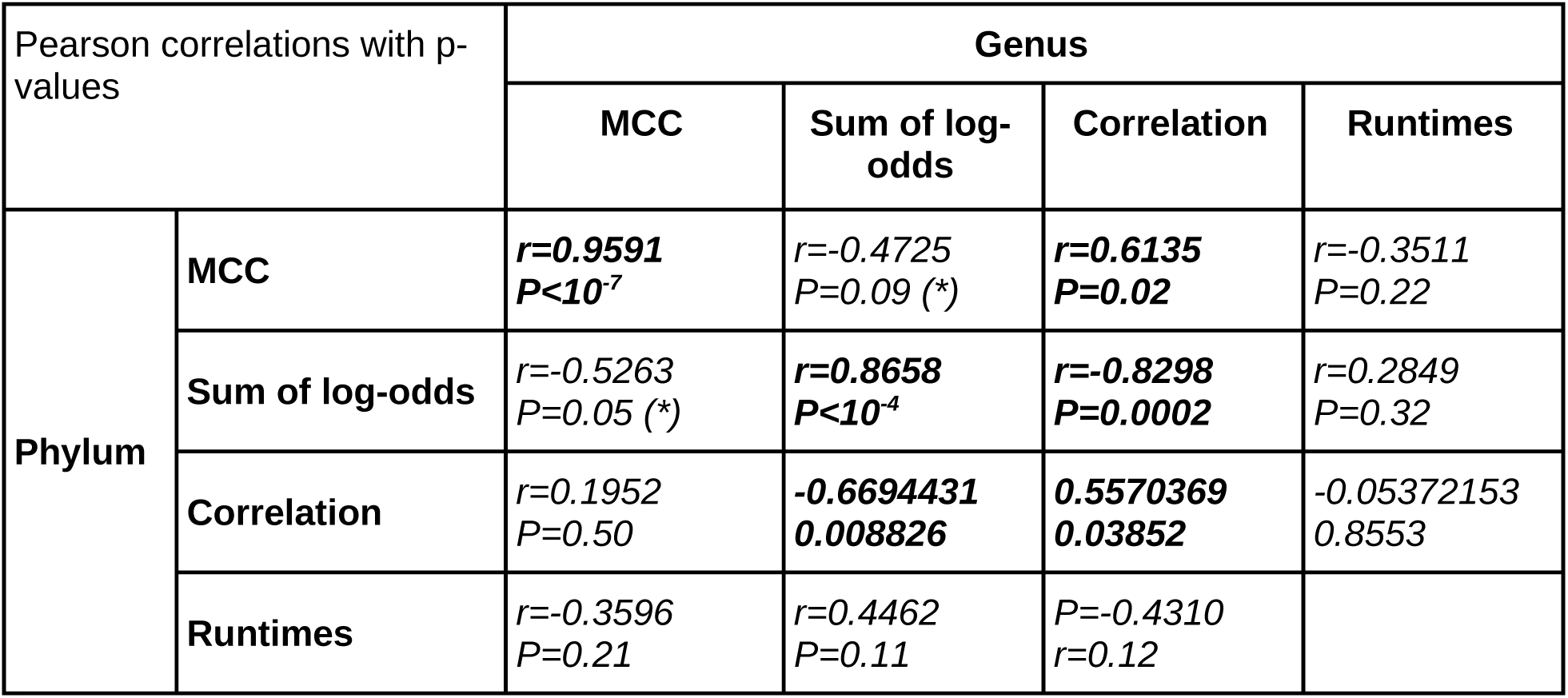
*Phylum level performance metrics for the individual methods. Average numbers for the simulated data sets are given. The metrics are true positives (TP), false positives (FP), true negatives (TN) and false negatives (FN) as well as sensitivity (SEN), specificity (SPEC), positive predictive value (PPV), negative predictive value (NPV) and Matthew’s Correlation Coefficient (MCC).*

**Table 3:**
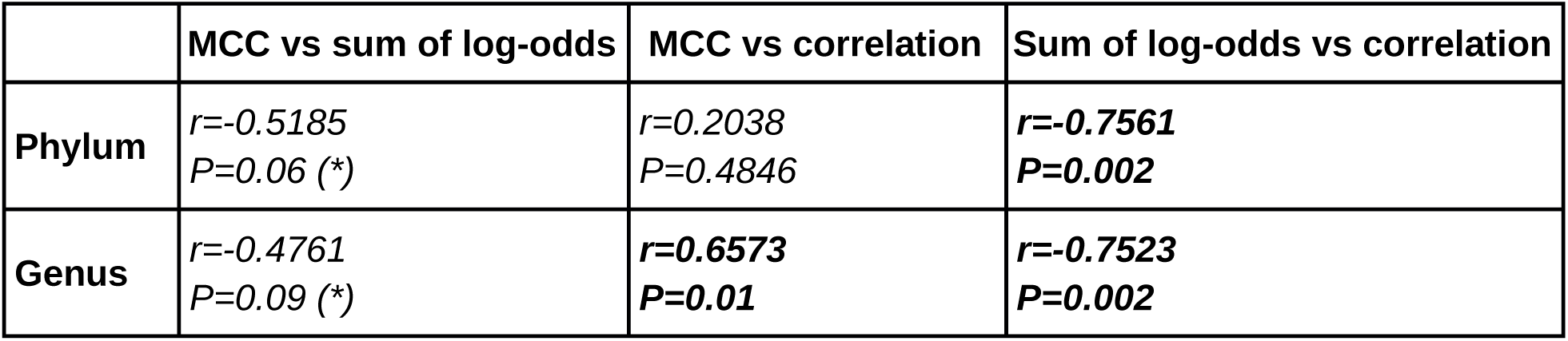
Pearson correlations between different quality metrics at the phylum and genus level. The metrics are Matthew’s Correlation Coefficient (MCC), the sum of log-odds scores between predicted and known proportions, and the Pearson correlation between all the predicted and known relative abundances. The run times are the same for both genus and phylum level. Significant correlations (P<0.05) are highlighted with bold. Marginally significant correlations (P<0.1) are indicated with asterisks (*).

**Table 4:**
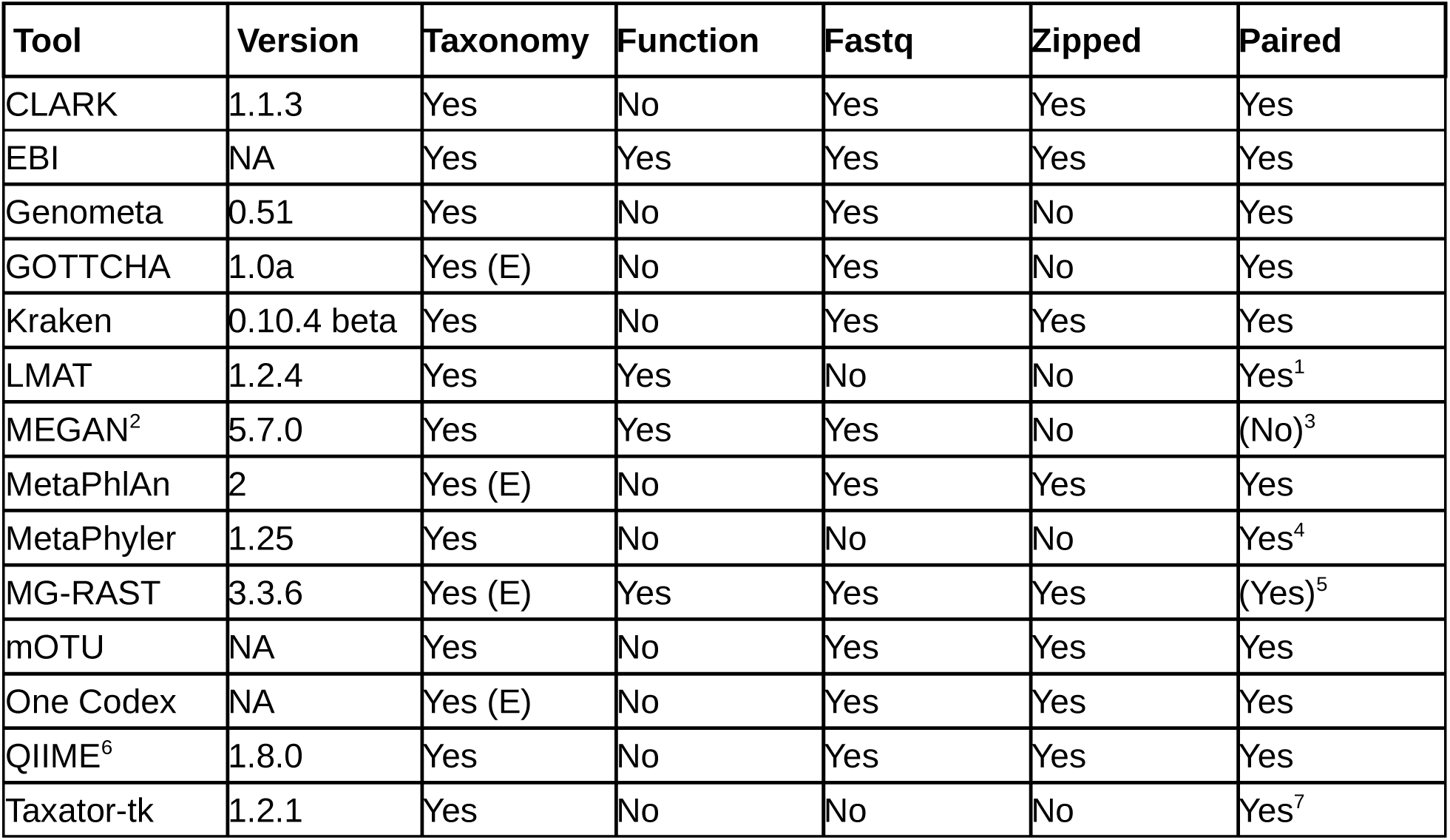
Comparison between different quality metrics at the level of either phylum or genus. Same metrics and notation as in Table 3.

Log-odds scores were used to investigate the differences between the known and predicted relative abundance of phyla and genera. Scores close to 0 indicate predictions close to the known proportions, a negative score means the program predicted too few, and a positive score means the program predicted too many. On average the dominant phyla (Proteobacteria, Actinobacteria, Acidobacteria) lie closer to 0 as it is less likely to predict a fold change in abundance for these groups (Supplementary Figure 2). Conversely, the rare phyla show the largest fluctuations by this metric. The simulated Spirochaetes are underrepresented by all tools, most likely because the genomes that were artificially evolved are difficult to confidently map to real genome sequences in the databases.

**Figure 2:**
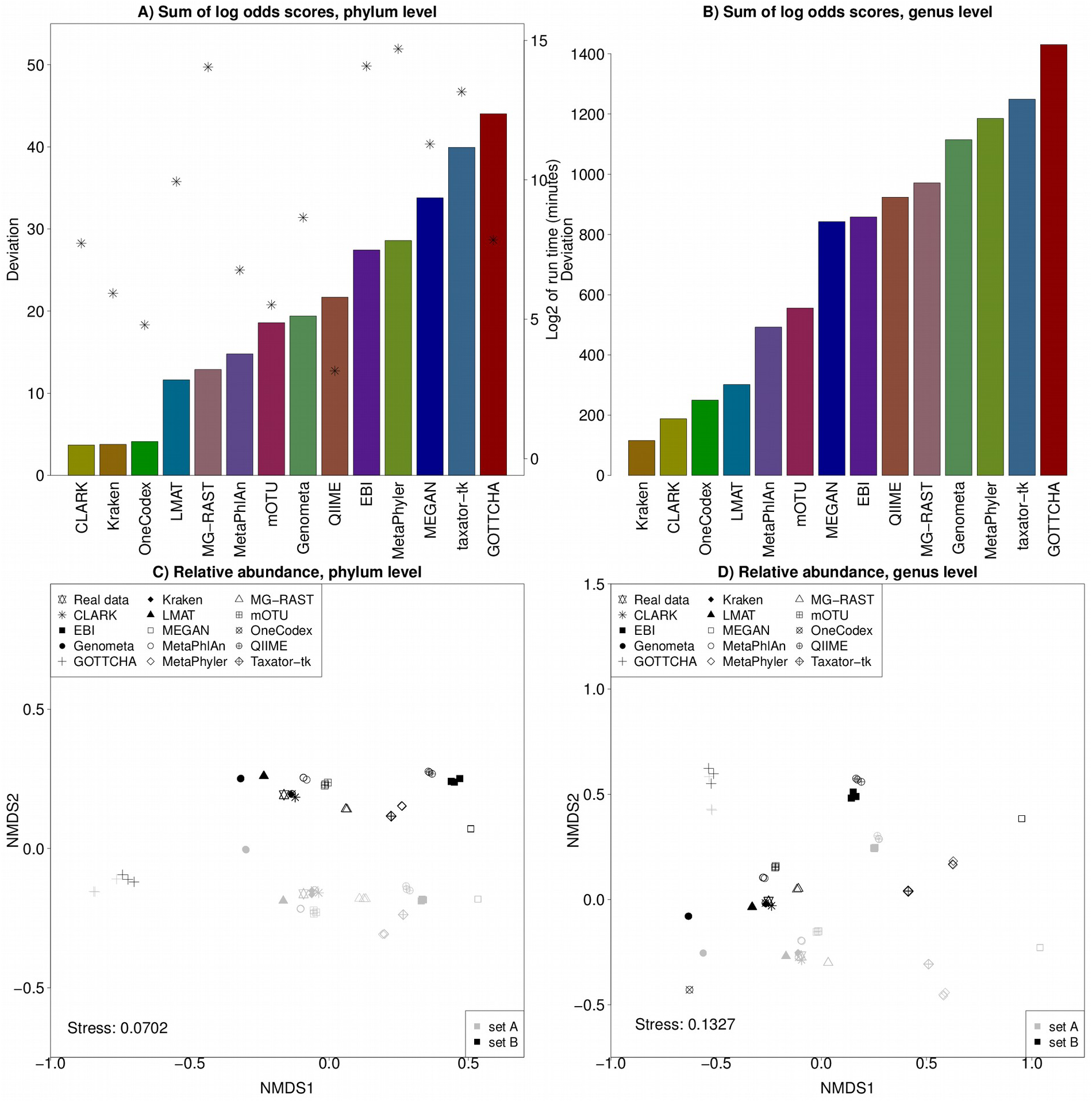
*Analysis of performance at the level of phylum (left) and genus (right). **A** and **B**: Sum of absolute log-odds scores at the phylum (A) or genus (B) level for each tool (bars) and log_2_ of run time in minutes (asterisks, ^*^). Sum of log-odds scores indicate the overall performance in terms of deviation from the known proportions. A low sum indicates a high accuracy. **C** and **D**: NMDS plot of relative abundances at the level of phylum (C) and genus (D) for the known and predicted communities in replicates. Eukaryotes are not included. Metagenomes in set A are gray, and metagenomes in set B are black. The known communities are shown with a star.*

Although all tools generate relative abundances that differ from the actual abundances, the degree to which they diverge is highly variable. In Figure 2A, the sum of absolute log-odds scores is shown for each tool at the phylum level, giving an overall indication of how much the results from each tool tend to diverge from the correct proportions (a small sum is preferable). Also shown is log_2_ of the run time in minutes, and there is no correlation between run times and sum of log-odds scores (Pearson correlation; *r* = 0.28, *P* = 0.32). In Figure 2B, the absolute sum of log-odds are shown at the genus level. As expected, the absolute deviation between the predicted and real relative abundances vary more at the genus level than at the phylum level, yielding larger sums of log-odds. Overall, the relative performance of the methods remains almost the same, with the four methods showing the lowest sum of log-odds being the same as before (Kraken, CLARK, OneCodex and LMAT). Some methods show a decreased performance at the genus level including MG-RAST (moving from position 5 to position 10) and Genometa (from position 8 to position 11), whereas MEGAN shows an improved relative performance (from position 12 to position 7). The sum of log-odds show a strong Pearson correlation (r = 0.87, *P* < 10^−4^) between phylum and genus level assignments.

The MCC and sum of log-odds are both measures of how well the predictions match the known abundances. However, the metrics are not significantly correlated at the phylum level (*r* = -0.52, P = 0.06) or the genus level (*r* = -0.48, P = 0.09). The reason for this is most likely that, as mentioned above, the sensitivity scores are biased for some methods. Indeed, if these methods are left out, the remaining methods show significant correlations at both the phylum level (*r* = -0.73, P = 0.03) and genus level (r = -0.81, P = 0.008).

The sum of log-odds should not be considered alone as they can be strongly affected by a single prediction being wrong (e.g. Taxator-tk on Gemmatimonadetes). Likewise, rare phyla (e.g. Elusimicrobia with only 0.44% of the reads) can have a disproportionate impact on the sum because a small absolute error can have a large relative impact on the log-odds. Therefore, Pearson correlations between predicted and known proportions were also calculated for each tool (Table 1). For each tool, only phyla predicted by that tool were included in the calculation. Although all the predictions at the phylum level correlate with the real abundances (all P<0.01), the correlation coefficients vary from *r* = 0.74 (EBI) to *r* = 0.99 (CLARK, Kraken). GOTTCHA is an outlier with very weak correlation (*r* = 0.18, *P* = 0.34). At the genus level, all correlations between predicted and known proportions are highly significant (all P<<0.001), but as was true for the phylum level, the strength of the correlation varies between methods, with both CLARK and Kraken having *r* = 0.96, and MEGAN showing the lowest correlation (*r* = 0.50).

The performance of the tools, as measured by correlations between predicted and known proportions, correlate significantly between the phylum and genus level (*r* = 0.56, *P* = 0.04) showing overall consistent performance. However, the performance does not correlate with run times at the phylum level (*r* = -0.05, *P* = 0.86) or the genus level (*r* = -0.43, *P* = 0.12). As expected, there is a negative correlation between sum of log-odds and average Pearson correlations at both the phylum level (*r* = -0.76, *P* = 0.002) and the genus level (*r* = -0.75, *P* = 0.002), indicating that a prediction that correlates well with the known composition also has a low sum of log-odds scores.

To visualize the degree of overall similarity between the predicted and the real communities, a non-metric multidimensional scaling (NMDS) ordination of the relative abundances was generated at the phylum level (Figure 2C) and the genus level (Figure 2D). There are three replicates of each set with identical proportions. Thus, each tool has three predictions per set, and ideally the corresponding symbols should be on top of each other.

In Figure 2C & D, there is a clear separation between sets A and B. Furthermore, most tools predict almost identical relative abundances for the three replicates in each set, placing the three individual points almost on top of each other (the largest variation is seen for EBI, GOTTCHA and QIIME). However, the similarity between predicted and known relative abundances (as measured by the distance between a set of predictions and the known proportions) varies between tools. The results from CLARK, Kraken and OneCodex are the closest to the actual points in both sets of metagenomes followed by Genometa, LMAT, MetaPhlAn and mOTU. The results from EBI, GOTTCHA and MEGAN are the most divergent, and the remaining tools produce results between the two extremes. The NMDS plot shows a similar pattern for the genus level performance. A cluster of programs predict relative abundances very close to the known distribution (Kraken, CLARK, OneCodex, LMAT), with another group of programs being close as well (MetaPhlan, mOTU, MG-RAST, Genometa).

### Functional analysis

Inferring function from metagenomes is much more challenging than inferring taxonomy. However, the extra information not only makes better use of the shotgun data but also adds a new and more ecologically relevant layer of information to the study. The tools LMAT, MG-RAST, MEGAN, QIIME/PICRUSt and the EBI metagenomics server all analyze the functional capacity of a metagenome by analyzing protein coding genes (with the exception of PICRUSt which infers protein-coding gene content from taxonomic profiles generated by QIIME).

The test datasets were created with differences in the relative abundance of cyanobacteria (photosynthesis; more abundant in set A), *Bradyrhizobium* and *Rhizobium* (nitrogen fixation; more abundant in set A), and known pathogens (more abundant in set B). Using the known shifts in taxa as a proxy for the expected functional shifts makes it possible to compare the predicted differences to what the expected change should be. Although the magnitude might differ due to e.g. the number of genes associated with a specific functional role, the genome size and the number of species sampled, the direction should be the same.

MEGAN, MG-RAST and LMAT all use the SEED hierarchy for functional annotation, and we used the same subset of SEED subsystems for these analyses although some subsystems were not predicted by all tools. PICRUSt uses KEGG, and the EBI webserver uses Gene Ontology (see Supplementary Material for the exact categories used, and Supplementary Table 3 for the predicted proportions). If more than one category was used for a function the average log fold change is reported. The shifts in functional categories are shown in Figure 3.

**Figure 3:**
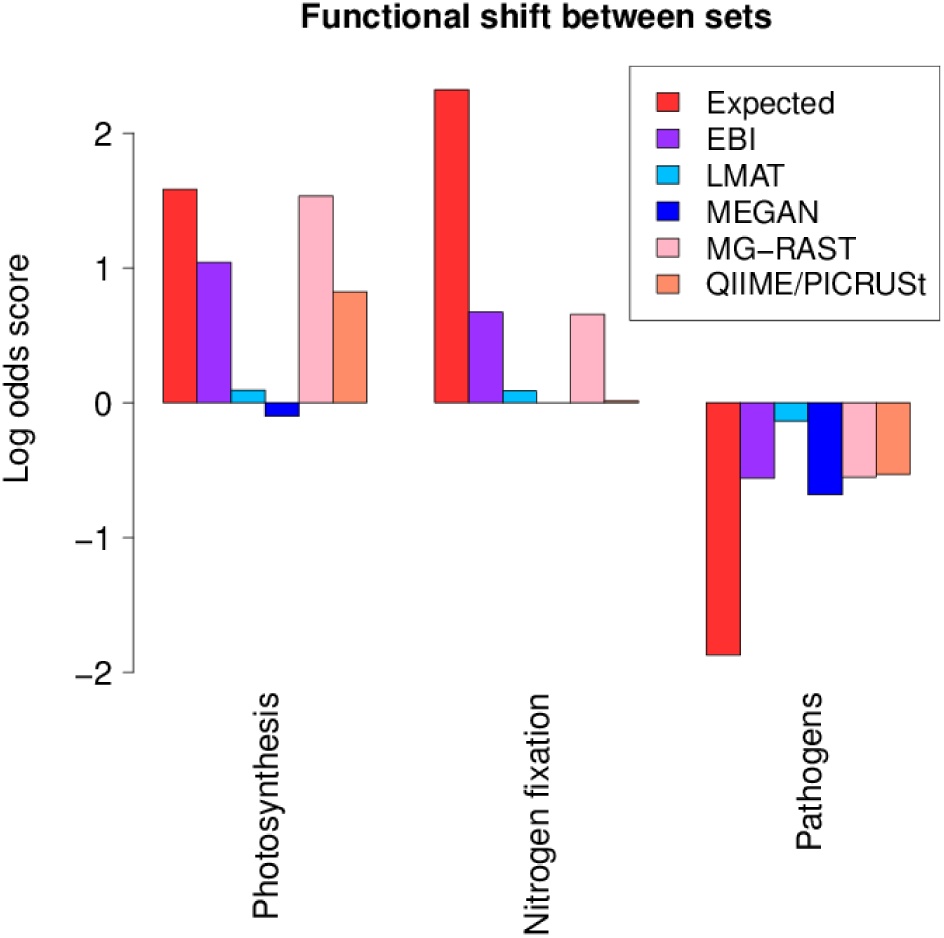
*Shifts in relative abundance of the three functional categories (or set of categories) that vary between set A and set B for the tools that analyze the functional capacity of metagenomes. A positive log-odds score means an increase in set A relative to set B, and a negative log-odds score means a decrease in set A relative to set B.*

Only MG-RAST and the EBI webserver capture the overall pattern of functional changes in all three functional roles. The direction is correct in all cases but the magnitude of the shifts differ. The closest match is in the photosynthesis category with MG-RAST almost mirroring the expected shift. For both nitrogen fixation and pathogens the shift is much less than what would be expected based on read counts alone. For the pathogens, one explanation is that a number of categories were included, some of which are not exclusive to pathogens. Thus, we would not expect a clean “pathogens” signal.

Although QIIME/PICRUSt rely on 16S rRNA genes to infer functional content, they predict a shift in the expected direction in both the photosynthesis and pathogens categories. There was no nitrogen fixation category predicted so the ‘Nitrogen Metabolism’ was used showing no difference between the two sets of metagenomes. The strategy employed by PICRUSt differs fundamentally from the other tools used so the performance is worth noting.

MEGAN shows a shift close to zero for photosynthesis and assigns no reads to the nitrogen fixation subsystem. Only for the pathogen-associated categories does the prediction match the expected change, showing the largest fold change of the tested tools. LMAT predicts minor shifts in all three categories, all in the right direction but too close to zero to be useful for most downstream analyses.

Given that the “pathogens” category contains a number of characteristics for which we would expect a mix of pathogens and non-pathogens (e.g. motility), it is interesting to see that most tools predict the expected shift between the sets. Moreover, the tools seem to agree on the magnitude of the shift as well. This is an important result as shotgun metagenomes are increasingly used to infer microbial community traits^30^, which are not necessarily linked to a single functional category. The photosynthesis category shows the best match between both direction and magnitude of the shift for three of the tools. For nitrogen fixation, only two tools (the EBI webserver and MG-RAST) predicted the expected shift.

## Discussion

This paper presents the first large independent benchmark of metagenome analysis tools using complex, realistic simulated data sets with replicates. The results show that the run time of individual tools varies by many orders of magnitude, and that the fraction of reads analyzed varies dramatically (from <1% to >70%). Although no tool predicts the actual relative abundances of bacterial phyla correctly (all tools differed significantly from the actual distributions), some predictions are much more similar to the correct answer than others. Interestingly, there is no correlation between the number of reads used, quality of the result and run time. Indeed, some of the best tools in terms of both similarity to the correct answer and the fraction of reads used are CLARK and Kraken, and these are also among the fastest tools tested. Picking a single “best tool” is not straightforward, but the different performance metrics presented in this benchmark can help researchers decide based on their specific demands.

Most tools only analyze the taxonomic distribution and do not look at the potential functional differences in terms of protein-coding gene content. For the tools that were able to perform this type of analysis (LMAT, MEGAN, MG-RAST, QIIME/PICRUSt, EBI), the predictions differed starkly between the tools. The EBI and MG-RAST webservers both predicted the expected direction of change in all three functional categories. Interestingly, using QIIME and PICRUSt predicted two of the functional changes, although the data used is not what these methods were developed for. The remaining methods predicted one (MEGAN) or none (LMAT) of the functional shifts.

The field of metagenomics is changing rapidly, and there is still room for improvement in the development of analysis tools. For taxonomic assignment, there are now tools that we show to perform well on controlled but complex and realistic data sets. One area of further research could be to focus on broadening the taxonomic range to include at least some eukaryotes and viruses of interest. When it comes to functional capacity, there seems to still be room for improvement on existing tools. Most of the tools available that are able to look at the functional capacity of metagenomes are among the slowest tools tested, so it might be fruitful to infer other approaches - potentially inspired by the fastest tools for taxonomic assignment. More importantly, only two of the five tested methods predicted all the functional changes, whereas the others did not. It might be worth looking into the underlying assumptions behind the functional assignments of reads. Here, it seems that the approaches taken by the EBI webserver and MG-RAST perform the best, but it is also worth noting the performance of PICRUSt although it only considers the taxonomic predictions from QIIME. This approach seems to be a relevant area for further research.

## Methods

To make the benchmark unbiased and realistic, the data sets used for testing had to mimic the complexities - both in terms of number of taxa present, observed shifts in the abundance of taxa, sequencing errors, and unknown reads that real shotgun metagenomes produce. A suite of analysis tools were then used to analyze the same data sets using the recommended settings for each tool, and the output from each tool was parsed in order to assess performance. In the following, details on the design of data sets, selection of tools, and analyses are presented.

### Creating test data sets

Designing the data sets was the most crucial part of the analysis. To ensure that the assessment was realistic and fair, we created data sets that closely mimic the complexity, size and characteristics of real data. Furthermore, to test whether the tools could distinguish between metagenomes from different communities we generated two sets - set A and set B - with different compositions. For both set A and set B, we generated three replicates. All replicates from a set have the same relative abundance of each phyla, but the actual genomes sampled as well as the positions within the genomes were randomized. Therefore, the benchmark was performed on 2 sets of metagenomes (set A and set B) with different characteristics (see below), each set contains 3 groups of simulated reads for a total of 6 simulated metagenomes.

We have decided to focus on two levels of taxonomic resolution in this benchmark - phylum and genus. From a practical perspective, comparing the phylum level predictions between tools is much easier and less prone to differences in e.g. naming conventions. Also, in a real metagenome study, many sequences will not match specific species due to biases in the databases and undersampling of large regions of the phylogeny. To investigate the performance at low and high resolution, we analyze the performance at both the phylum and genus levels, thus avoiding problems with naming of strains and species while still being able to benchmark how well the individual tools are able to assign reads to groups of varying specificity.

To capture the intricacies of next-generation sequencing, the data sets were based on real sequencing results obtained from pooling 6 soil DNA samples on a single HiSeq 2000 lane and generating 2x100 PE shotgun reads (unpublished, manuscript in preparation). The real sequence reads were discarded for this benchmark, but an error profile was calculated for each of the 6 libraries using ART^31^ based on the reported quality scores. Using ART and the error profiles, read pairs were simulated (read length 100, mean insert size 500, standard deviation 25) from both real, simulated and shuffled genomes (see below). The resulting 6 metagenomes contain between 27 and 37 million read pairs.

To make comparison between tools possible, the relative abundance of individual phyla was controlled by sampling read pairs from sequenced genomes in well defined proportions (see Supplementary Table 1 for the exact proportions and number of reads, and Supplementary Table 2 for the list of genomes used in the different categories). Some phyla were included in equal proportions in all data sets, while others varied either subtly or more substantially between sets A and B. Note that eukaryotes are treated as a single “phylum” although the chromosomes used come from various phyla (including e.g. Arabidopsis, chicken and human). By sampling reads at the genome level, we can also calculate the relative abundance of individual phyla as well as for use in the comparison.

In many published metagenome studies, a large number of the reads can not be given a taxonomic assignment, the most likely reason being that many of sequences are from unknown organisms that are not present in the reference databases being used. To mimic this pool of unknown reads, all the simulated metagenomes contain 20% shuffled reads obtained by shuffling a set of 110 genomes using the shuffle program from the HMMER package^32^ (using parameters “-d -w 500” to use a local window and preserve dinucleotide distribution) and sampling reads from these shuffled “genomes”. These reads served as a negative control as the genomes of origin were not included in the simulated communities.

Since taxonomic assignment is based on sequenced genomes, there will be an inherent bias towards classification of reads from groups that have been sampled frequently (e.g. human pathogens) whereas genomes from less studied branches of the tree of life (e.g. Archaea) will often not be assigned. Furthermore, reads from an unsequenced relative of a known species might be easy or hard to assign depending on the evolutionary distance between the two. To test how the different tools perform on sequences that diverge from the available genomes but are not shuffled as in the above test, we used Rose version 1.3^33^ to simulate evolution and generate simulated relatives of the Spirochaete, *Leptospira interrogans* (EMBL ID AE016823). In total, a set of 32 genomes was generated containing 8 genomes with either little, medium, mixed or high divergence. No other Spirochaete genomes were included in the data sets. Since these genomes are not random but simulated using an evolutionary model, the reads should be assigned to the correct clade.

Some tools are able to not only determine community composition from a metagenome but can also infer “functional capacity” by analyzing the predicted protein coding sequences. To test the latter, the two sets were designed with functional differences: 1) the proteobacteria contain different proportions of known pathogens, 2) the genera Rhizobium and Bradyrhizobium, which are capable of nitrogen fixation, vary, and 3) the relative abundance of cyanobacteria (photosynthesizers) differ between the two sets. The different contributions of these groups to set A and set B should be reflected in both the relative abundance of phyla, genera and functional categories.

To evaluate the performance of each tool, no single metric was used as this is a complex problem where multiple factors should be considered. Instead, we investigated a number of performance metrics comparing the predicted to the known community structure, or looking at the specific performance of a tool:

- **Run time:** For the end user, the time spent analyzing data can be a significant bottleneck. We used the ‘GNU time’ (version 1.7) function to determine the cpu time spent on each data set, taking into account the use of multiple cores etc. A user can then assess if the time needed for a given tool to finish is realistic, and if the extra time required for some tools yields a proportional increase in the quality of the prediction.
- **Ease of use:** For the end user, how easy it is to use a tool can be a decisive factor. We did not specifically evaluate how user friendly a tool is, but we list some key things to consider: Can you use the zipped, paired-end Fastq files directly? Do you have to unzip the files? Do you have to convert the Fastq files to Fasta files? Does the tool utilize paired-end information? The number of steps involved varies between tools. In Section 2 of the supplementary material, we list the commands used in the analyses so the reader can judge how easy it is to run a tool.
- **Information provided:** Most tools only provide information on the community composition, while some provide functional information as well.
- **Reads mapped:** Since some tools use a limited set of marker genes for analyses, whereas others use much broader databases, the fraction of reads mapped in itself might not be useful. However, in case two tools both use the same or similar databases, it can be of interest to know how large a fraction of the reads each tool was able to analyze.
- **Shuffled reads mapped:** As a measure of specificity, it is of interest to know how many of the shuffled reads (generated from shuffled genomes) each tool maps to a phylum since this can reflect how often real reads might be wrongly assigned as well.
- **Non-existing phyla:** As we know exactly which phyla have been included in the data sets, we can also test how often a tool predicts the presence of a phylum that is not included in the data.
- **Divergence from real distribution:** Each predicted relative abundance of a phylum was compared to the actual abundance, and the overall divergence for each tool was evaluated.
- **Correlation with known community composition:** By comparing the relative abundance of phyla generated by each tool to the known composition, Pearson correlation coefficients were calculated giving the overall relationship between prediction and the real abundances.
- **Multivariate analysis:** The overall agreement between tools, and between tools and the real distributions, were visualized with non-metric multidimensional scaling (NMDS) plots which group similar predictions together and give a broad overview of agreements and disagreements.
- **Sensitivity, specificity, PPV, NPV and MCC:** Since the provenance of each read is known, we can also calculate the number of true positives, false positives, true negatives and false negatives for each tool and use these numbers to calculate sensitivity, specificity, positive predictive value (PPV), negative predictive value (NPV) and Matthew’s correlation coefficient (MCC).

In combination, these different metrics will help the end user decide which tool is best suited for their particular needs.

### Selection of tools

There are many tools available for analyzing metagenomes. For the present analysis, the following criteria were used to include tools:

- **Availability:** The tool should be freely available either as download or webserver.
- **Usability:** The tool should have a proper manual, readme file or help function describing how to use it. In case of problems, the respective authors were contacted.
- **Adoption:** The tool should be widely used, or show potential of being widely adopted in the future.

Any selection will by necessity exclude tools that some researchers would find useful. However, it is infeasible to test all available tools, and using the above criteria gives a clear way of narrowing down the options. Some tools were considered but excluded due to lack of support, lack of details on how to use the tool, or nonfunctioning webservers.

The following 14 tools were included in the analysis (also see Table 5):

**Table 5:**
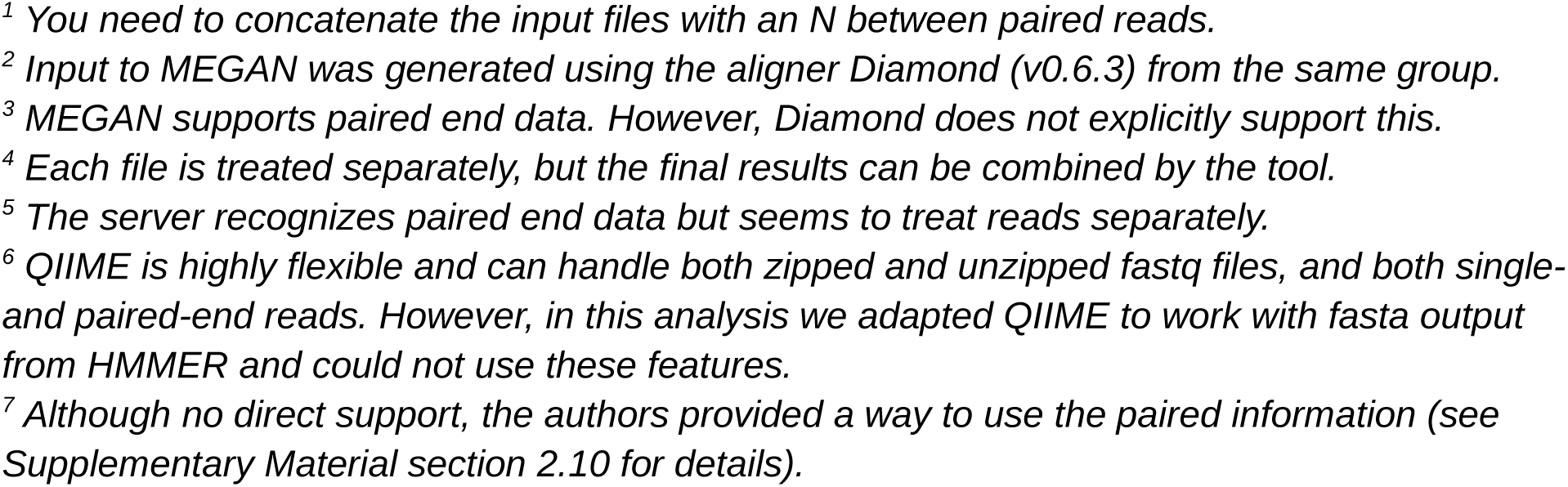
*The metagenome analysis tools included in this benchmark. For each tool, it is shown if it does taxonomic analysis (tools that can also infer Eukaryotic taxa are noted with an “(E)”) and/or functional analysis, and whether it can analyze Fastq files directly, if you can use zipped input files, and if it utilizes paired end information.*

- **CLARK^16^**: CLARK uses a *k*-mer approach where all the common *k*-mers between the targets in the database (e.g. a collection of all bacterial genomes) are removed. This gives a set of genomic regions that uniquely describes each target. A read is assigned to the target with which it shares the highest number of *k*-mers. In the most accurate mode, CLARK uses the full database of targets and assigns a confidence score to the assignment.
- **EBI metagenomics webserver** ^34^: Hosted by the European Molecular Biology Laboratory-European Bioinformatics Institute (EMBL-EBI), the webserver ties in with the European Nucleotide Archive (ENA) for data storage and demands a minimum of metadata. The raw reads are cleaned, trimmed using sff-trim from BioPython ^35^ and trimmomatic ^36^, clustered to remove duplicates using UCLUST ^37^, and repeats are masked using RepeatMasker ^38^. Paired end reads are joined using SeqPrep ^39^. Potential rRNA genes are found using rRNAselector ^40^, and the taxonomy is predicted from these using QIIME ^28^ against RDP ^41^ and GreenGenes ^42^. Protein coding genes are predicted using FragGeneScan ^43^, and the functional annotations are added using InterProScan ^44^ against a set of databases (including Gene3D ^45^, Pfam ^46^ and PROSITE ^47^), and each predicted gene is associated with a set of Gene Ontology terms ^48^.
- **Genometa** ^17^: Genometa is a GUI program building on the Integrated Genome Browser (IGB) ^49^ and using the Bowtie mapper ^50^. Reads are mapped to a custom genome database and results can be exported from the GUI program for downstream analysis. The most recent database was used (April 2012) together with Bowtie v1.1.0 which is the only option in the program.
- **GOTTCHA**^18^: GOTTCHA aims at limiting the number of false positives in their predictions by only focusing on genomic regions that are unique to each reference. These regions are found using a combination of empirical data on coverage and machine learning approaches. GOTTCHA first trims the reads based on qualities followed by fragmentation to obtain a uniform read length. A read is split on every low quality base (Q<20) and then divided into all non-overlapping 30-mers. Matching is done at the species level using exact matches with BWA^51^.
- **Kraken** ^19^: Kraken classifies reads by breaking each into overlapping k-mers. Each k-mer is mapped to the lowest common ancestor of the genomes containing that k-mer in a precomputed database. For each read, a classification tree is found by pruning the taxonomy and only retaining taxa (including ancestors) associated with k-mers in that read. Each node is weighted by the number of k-mers mapped to the node, and the path from root to leaf with the highest sum of weights is used to classify the read.
- **LMAT** ^20^: LMAT works by first generating a searchable database of k-mers from a large collection of genomes. For each k-mer, the lowest common ancestor in the taxonomy tree is calculated. A smaller “marker library” (kML) of the most informative k-mers is generated by separating k-mers into disjoint sets and discarding all sets with fewer than 1000 k-mers, and all k-mers where the lowest common ancestor is above the family level. When assigning taxonomy to a read, the information for each k-mer in the read is extracted from the library, and the path from the highest scoring node to its lowest common ancestor is created. This path is pruned by each conflicting assignment until the score drops below the threshold. Function is assigned in a similar fashion using a customized library.
- **MEGAN** ^22^: MEGAN is a GUI (Graphical User Interface) program aimed at analyzing a set of reads that have been mapped to a sequence database. The mapped reads are assigned a taxonomic label by finding the “lowest common ancestor” in the NCBI taxonomy based on sequence similarity. Functional analysis can be performed using different databases. Here, we use the SEED hierarchy ^52^. MEGAN assigns function to a read by mapping it to the best matching gene with a known function in the hierarchy. The initial mapping is performed using the RefSeq database ver. 66 ^53^ using the Diamond aligner ^54^ from the same group as MEGAN.
- **MetaPhlAn** ^25^: MetaPhlAn uses a set of around 1 million markers for taxonomic assignment and, thus is not expected to map all reads. The marker set is based on clade-specific sequences, where a clade can be as specific as a species or as broad as a phylum, and marker sequences have to be strongly conserved within a clade without being locally similar to sequences outside the clade. Taxonomic assignment is accomplished by mapping all reads against the marker set using Bowtie2 ^55^.
- **MetaPhyler** ^26^: MetaPhyler relies on a custom database of 31 marker genes for taxonomic assignment and is therefore not expected to map all reads in a data set. The marker set consists of mostly universal single-copy genes and Metaphyler has a specific classifier for each ^56^. MetaPhyler uses BLASTX ^57^ to assign reads to marker genes and uses the bit score (adjusted for alignment (HSP) length and marker gene) to assign taxonomy from genus to phylum level.
- **MG-RAST** ^23^: The MG-RAST webserver allows the user to upload metagenome data sets for analysis. The server offers an easy to use pipeline that performs both taxonomic and functional analyses using custom databases M5nr ^58^ for proteins and M5rna (combining SILVA ^59^, GreenGenes ^42^ and RDP ^41^) for rRNA analysis. Gene calling is performed on the reads using FragGenescan ^43^ and predicted proteins are clustered using UCLUST ^37^. BLAT ^60^ is used for calculating similarities for representatives from each cluster. Potential rRNA reads are found using BLAT against a reduced version of the SILVA database, reads are clustered using UCLUST, and BLAT is used against the M5rna database.
- **mOTU** ^27^: mOTU uses a set of 10 universal single-copy marker genes to assign taxonomic information to reads. These marker genes were chosen from a larger set of 40 genes as the ones that performed best in an internal benchmark by the authors of mOTU. A large number of genomes and metagenomes were scanned for these marker genes, and the results clustered into “metagenomic operational taxonomic units” (mOTUS). Taxonomic assignment of reads is performed by mapping them against this mOTU database.
- **One Codex^61^**: One Codex is a web platform (free for academic use) that uses a *k*-mer approach to compare uploaded FASTQ or FASTA files to their in-house index of microbial genomes. Exact *k*-mer matches are used to find the most specific taxonomic assignment for each read, using a lowest common ancestor approach. The output presents the results with a graphical overview of the sample composition, a break-down into high, medium and low abundance species, and direct access to read level classifications.
- **QIIME** ^28^: It is important to note that QIIME is a software package aimed at analyzing amplicon data (e.g. SSU rRNA sequences) and is thus not designed for the analysis of shotgun data sets such as the ones used in this paper. However, since QIIME is widely adopted and highly flexible, we found it interesting to test how it might be used on this type of data. Using HMMER (ver 3.1b1) ^32^ and rRNA alignments from Rfam (ver 12.0) ^62^ potential reads from 16S rRNA genes were found. This type of data differs from what QIIME is designed for, because it consists of short, random segments that cover different parts of the rRNA gene and that therefore differ in their taxonomic informativeness. It is therefore not the optimal input. QIIME was used to pick open-reference OTUs from Greengenes 13_8 ^42^ using UCLUST ^37^ and representative sequences were picked. Taxonomy was assigned using UCLUST. A functional analysis of the metagenomes is performed using PICRUSt ^63^, a tool that predicts the functional profile of microbial community. The gene content of each organism in a phylogeny (here GreenGenes) is precomputed. Next, the relative abundance of 16S rRNA genes is used to infer the gene content of the metagenome samples, taking into account the copy number of 16S genes. The final functional prediction is based on the KEGG Orthology ^64^.
- **Taxator-tk** ^24^: taxator-tk can use BLASTN ^57^ or LAST ^65^ to search with the reads against a local reference database. Overlapping local alignments between the read and a number of reference sequences are joined to form longer segments. Each segment is assigned a taxon ID, and a consensus taxonomic assignment is derived by assigning a weight to each segment based on similarities to the reference sequence.

Each tool was run using default settings and with the database recommended by the authors (i.e. a custom database in most cases). If needed, the authors of the individual tools were contacted to verify or troubleshoot the commands used. All authors of the tools used in this study have received a draft of the paper prior to publication in order to give their feedback. Command lines and settings for each tool can be found in the supplementary material.

Predicted abundances were extracted at the phylum level and analyzed as follows: all hits to Eukaryotes were grouped together as a single eukaryote ‘phylum’. Due to inconsistencies in naming, hits to Tenericutes and Firmicutes were summed and treated as Firmicutes. All hits to phyla not included in the simulated data sets were grouped and treated as “other”. R ^66^ was used to do all statistical analyses and to generate all the plots. The result from each tool was compared to the simulated abundances (excluding Eukaryotes for tools that only mapped to bacteria and archaea) with Pearson correlation coefficients.

The divergence between predicted and simulated abundances was calculated doing simple logodds scores for each method:

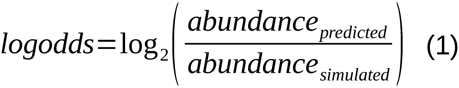

It should be noted that we only include log-odds for phyla or genera that were simulated in the data sets (thus, we always have abundance_simulated_>0) and which were predicted by the method being evaluated (thus, we always have abundance_predicted_>0). This avoids any issues with division by 0 or scores approaching infinity.

When summing these, the absolute values were used to measure total divergence irrespective of whether a tool over- or underrepresented specific phyla. To calculate sensitivity (SEN), specificity (SPEC), positive predictive value (PPV), negative predictive value (NPV) and Matthew’s Correlation Coefficient (MCC), we need to define true positives (TP), false positives (FP), true negatives (TN) and false negatives (FN). Since we know the provenance of each read, we can get FP as the number of shuffled reads that were mapped plus the number of real reads that were mapped to non-existing phyla. The number of TN is the number of shuffled reads that were not mapped. Similarly, by summing the number of real reads that were mapped, we get TP and from that FN follows as the number of real reads that did not map. We can then calculate the quality metrics:

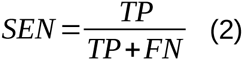

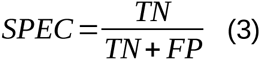

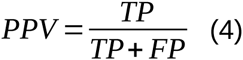

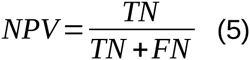

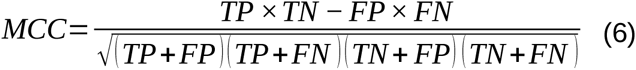

NMDS plots were generated using method metaMDS from the vegan ^67^ package with Bray-Curtis distances. For functional predictions, categories related to photosynthesis and nitrogen-fixation were assessed directly. For pathogens, a number of functional categories that have been shown to potentially be associated with pathogenicity were used (apart from virulence these include motility ^68^, sporulation ^69^ and quorum sensing ^70^).

## Author contributions

SL, KLA and PPG designed the experiment. SL created the data sets, ran the tools and analyzed the results. SL wrote the paper with input from KLA and PPG. All authors read and approved the final manuscript.

## Acknowledgements

SL, KLA and PPG would like to thank the authors of the different tools tested in this paper for their feedback. SL, KLA and PPG would also like to thank the many commenters on the preprint version for valuable feedback, in particular Thomas Sharpton (@tjsharpton), Paul Igor Costea (@CosteaPaul) and Genivaldo Silva (@meta_geni). SL, KLA and PPG are not in any way affiliated with any of the groups behind the software tested in this paper, and there is no conflict of interest. SL is supported by a Marie Curie International Outgoing Fellowship within the 7th European Community Framework Programme. KLA is supported by a postdoctoral fellowship from the Allan Wilson Centre for Molecular Ecology and Evolution. PPG is supported by a Rutherford Discovery Fellowship administered by the Royal Society of New Zealand

## Additional Information

The authors declare no competing financial interests.

---

Tables and Table legends:

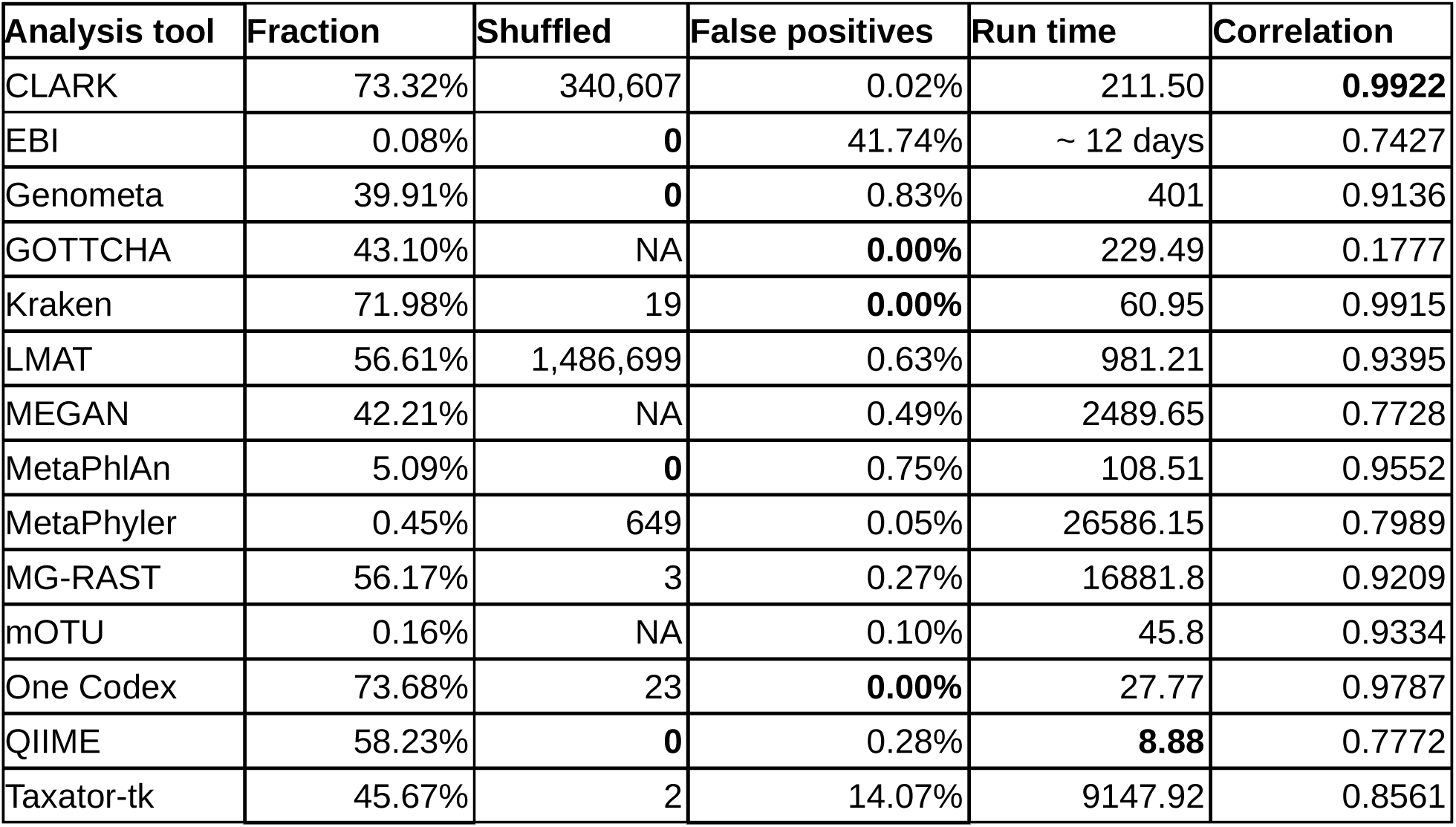

